# Nascent transcript analysis of glucocorticoid crosstalk with TNF defines primary and cooperative inflammatory repression

**DOI:** 10.1101/524975

**Authors:** Sarah K. Sasse, Margaret Gruca, Mary A. Allen, Vineela Kadiyala, Tengyao Song, Fabienne Gally, Arnav Gupta, Miles A. Pufall, Robin D. Dowell, Anthony N. Gerber

## Abstract

The glucocorticoid receptor (GR) binds to specific DNA sequences and directly induces transcription of anti-inflammatory genes that contribute to cytokine repression, frequently in cooperation with NF-kB. Whether inflammatory repression also occurs through local interactions between GR and inflammatory gene regulatory elements remains controversial. Here, using Global Run-on Sequencing (GRO-seq) in human airway epithelial cells, we show that glucocorticoid signaling represses transcription within 10 minutes. Many repressed regulatory regions reside within ‘hyper-ChIPable’ genomic regions that are subject to non-specific interactions with some antibodies. When this was accounted for, we determined that transcriptional repression occurs without local GR occupancy. Instead, widespread transcriptional induction through canonical GR binding sites is associated with reciprocal repression of distal TNF-regulated enhancers through a chromatin-dependent process, as evidenced by chromatin accessibility and enhancer-reporter assays. Simultaneously, transcriptional induction of key anti-inflammatory effectors is decoupled from primary repression through cooperation between GR and NF-kB at a subset of regulatory regions. Thus, glucocorticoids exert bimodal restraints on inflammation characterized by rapid primary transcriptional repression without local GR occupancy and secondary anti-inflammatory effects resulting from transcriptional cooperation between GR and NF-kB.

## INTRODUCTION

Glucocorticoids play a crucial role in normal physiology and are highly effective anti-inflammatory drugs with a remarkable range of clinical indications including asthma, rheumatoid arthritis, lupus, inflammatory bowel disease, psoriasis, and vasculitis, among others (Morand 2000; Barnes 2006; Gerber 2015; Kim et al. 2017). Glucocorticoids exert their potent biological and pharmacological effects through binding to the glucocorticoid receptor (GR), which causes GR to translocate to the nucleus and regulate gene expression through directly interacting with specific DNA sequences (Meijsing 2015; Sacta et al. 2016). Expression changes caused by glucocorticoids include gene induction and repression, with repression encompassing negative regulation of responses to inflammatory signals such as tumor necrosis factor (TNF) and lipopolysaccharide (LPS), including robust repression of cytokine expression (Rao et al. 2011; Uhlenhaut et al. 2013). Consequently, transcriptional repression is central to glucocorticoid-mediated anti-inflammatory effects (Clark and Belvisi 2012; Chinenov et al. 2013).

Pre-genomics experiments (La Baer and Yamamoto 1994), subsequently confirmed using deep sequencing-based approaches (Biddie et al. 2011; John et al. 2011), have established that inductive gene regulation by GR is typically nucleated through protein-DNA interactions between homodimeric GR and high affinity palindromic or semi-palindromic consensus GR binding sequences that are found in regulatory regions of glucocorticoid– induced genes (So et al. 2007). Mechanisms underpinning GR-mediated gene repression are less well understood. Although protein products resulting from GR-induced gene expression, such as TSC22D3 and DUSP1, are known to indirectly contribute to glucocorticoid-mediated transcriptional repression (Auphan et al. 1995; Ronchetti et al. 2015; Newton et al. 2017), direct repressive effects of GR on inflammatory transcription factors, such as NF-kB, have long been viewed as principally responsible for the potent repressive effects of glucocorticoids on cytokine expression (Cruz-Topete and Cidlowski 2015; Vandewalle et al. 2018). Such primary repressive effects have been variably attributed to protein-protein tethering of monomeric GR to DNA-associated inflammatory transcription factors, commonly referred to as transrepression (Ratman et al. 2013; De Bosscher et al. 2014), and also to protein-DNA interactions between GR and so-called negative glucocorticoid response elements (nGREs) found within regulatory regions for inflammatory genes (King et al. 2013). Both mechanisms are purported to ultimately result in GR-centered recruitment of repressive complexes and down-regulation of specific inflammatory genes.

Significant controversy has emerged regarding both of these putative repressive mechanisms. Enrichment for nGRE sequences within GR-occupied regions has not been evident on a genome-wide basis (Rao et al. 2011; Kadiyala et al. 2016; Oh et al. 2017). Similarly, repressive tethering interactions between GR and NF-kB have not been observed in ChIP-seq studies of GR occupancy in the setting of NF-kB activation (Uhlenhaut et al. 2013; Oh et al. 2017). Accordingly, the notion that GR-mediated repression is largely secondary, that is a result of GR-induced targets exerting repressive effects, has recently been suggested (Oh et al. 2017). However, analysis of gene expression in the presence of cycloheximide has indicated that protein synthesis is not required for at least partial glucocorticoid-based repression of selected inflammatory genes (King et al. 2013). Recent work has also characterized the structural basis for interactions between GR and nGREs (Hudson et al. 2013), suggesting that such interactions could at least theoretically occur. Thus, there is vigorous ongoing debate and discordant reports regarding the fundamental mechanisms that underpin GR-mediated gene repression (Oh et al. 2017; Sacta et al. 2018). A definitive answer to this question would have crucial implications for both understanding glucocorticoid-resistant inflammation and the development of improved anti-inflammatory therapies.

To address the mechanistic basis for glucocorticoid-mediated repression, we previously applied ChIP-seq to define occupancy of GR, the p65 subunit of NF-kB, and RNA polymerase II (RNAPII) in Beas-2B airway epithelial cells treated for one hour with the potent synthetic glucocorticoid, dexamethasone (dex), TNF, and both dex and TNF (Kadiyala et al. 2016). Whereas our studies revealed rapid and robust reduction in RNAPII occupancy across a host of pro-inflammatory genes when cells were treated with dex + TNF in comparison to TNF alone, our data did not confirm either a transrepressive or nGRE mechanism underlying these inhibitive effects on NF-kB. In contrast, we discovered that GR and NF-kB cooperate to enhance the expression of key anti-inflammatory genes, in a process mediated through consensus binding sequences for both transcription factors within individual enhancers. Targets regulated through this novel anti-inflammatory mechanism include *TNFAIP3* (frequently referred to as A20), *TNIP1*, *SERPINA1* (which encodes Alpha1-antitrypsin), and *NFKBIA*; our data further established that targets of GR-NF-kB cooperation are required for the full repressive effects of glucocorticoids on gene expression of various TNF-induced cytokines (Altonsy et al. 2014; Sasse et al. 2016). However, our studies lacked sufficient temporal resolution to determine definitively whether GR-mediated repression is effected solely as a secondary consequence of inductive gene regulation, including targets of GR-NF-kB cooperation such as TNFAIP3, or whether primary repression of NF-kB activity by GR also contributes to inflammatory repression by glucocorticoids.

Techniques that combine nuclear run-on approaches with deep sequencing (for example, Global Run-on Sequencing or GRO-seq) have been extremely useful in defining enhancer activity and nascent transcriptional changes associated with signal transduction (Hah et al. 2011; Allen et al. 2014). Thus, to address mechanisms through which glucocorticoids exert repressive effects on TNF signaling, we performed GRO-seq on Beas-2B cells treated with dex, TNF, and dex + TNF for 10 or 30 minutes. Through integrating our GRO-seq results with ChIP-seq data, chromatin accessibility assays, and enhancer activity analysis, we aimed to determine definitively whether GR exerts classically described direct repressive effects on NF-kB, or instead whether inflammatory repression can be fully attributed to other mechanisms.

## RESULTS

### GRO-seq defines rapid effects of dex and TNF on gene transcription and identifies new anti-inflammatory targets of GR signaling

To determine mechanisms and targets through which ligand-activated GR represses gene expression, we performed GRO-seq on Beas-2B airway epithelial cells treated for 10 and 30 minutes with vehicle (ethanol), TNF, dex or TNF and dex in combination. Airway epithelial cells were selected for these experiments as they are a target of glucocorticoid treatment in highly prevalent diseases such as asthma and chronic obstructive pulmonary disease (Gerber 2015). Differential transcription analysis (using an adjusted p-value p_adj_ < 0.05) of both TNF and dex treatments relative to vehicle revealed that numerous genes show a robust transcriptional response after just ten minutes (Fig. 1). At this early time point, the effects of TNF were exclusively inductive (Fig. 1A). Similarly, the majority of dex-regulated loci at this time point were induced (Fig. 1B), although 12 genes (e.g. *KRT17*, Fig. 1B) exhibited statistically significant reduction in transcription after 10 minutes of dex treatment. Given estimated rates of mammalian RNAPII transcription, mRNA maturation and transport, and translation (Hargrove et al. 1991; Schwanhäusser et al. 2011; Cheng et al. 2016), both the 10 and 30 minute time points are likely too rapid for protein synthesis from newly transcribed genes to occur. Supporting this notion, western blot analysis of TNFAIP3, NFKBIA, and CEBPB, all of which are transcriptionally induced by GR (Supplemental File S1), did not demonstrate detectable increases in protein expression after 10 or 30 minutes of dex treatment (Supplemental Fig. S1). Thus, the GRO-seq data are consistent with a direct repressive effect of ligand-activated GR on baseline RNAPII activity at specific genomic regions. Moreover, using a non-adjusted p-value cutoff of < 0.05, over 200 genomic regions exhibited reduced transcription after 10 minutes of dex exposure, suggesting that primary repressive effects (i.e. not requiring a protein intermediate) are widespread.

**Figure 1.**
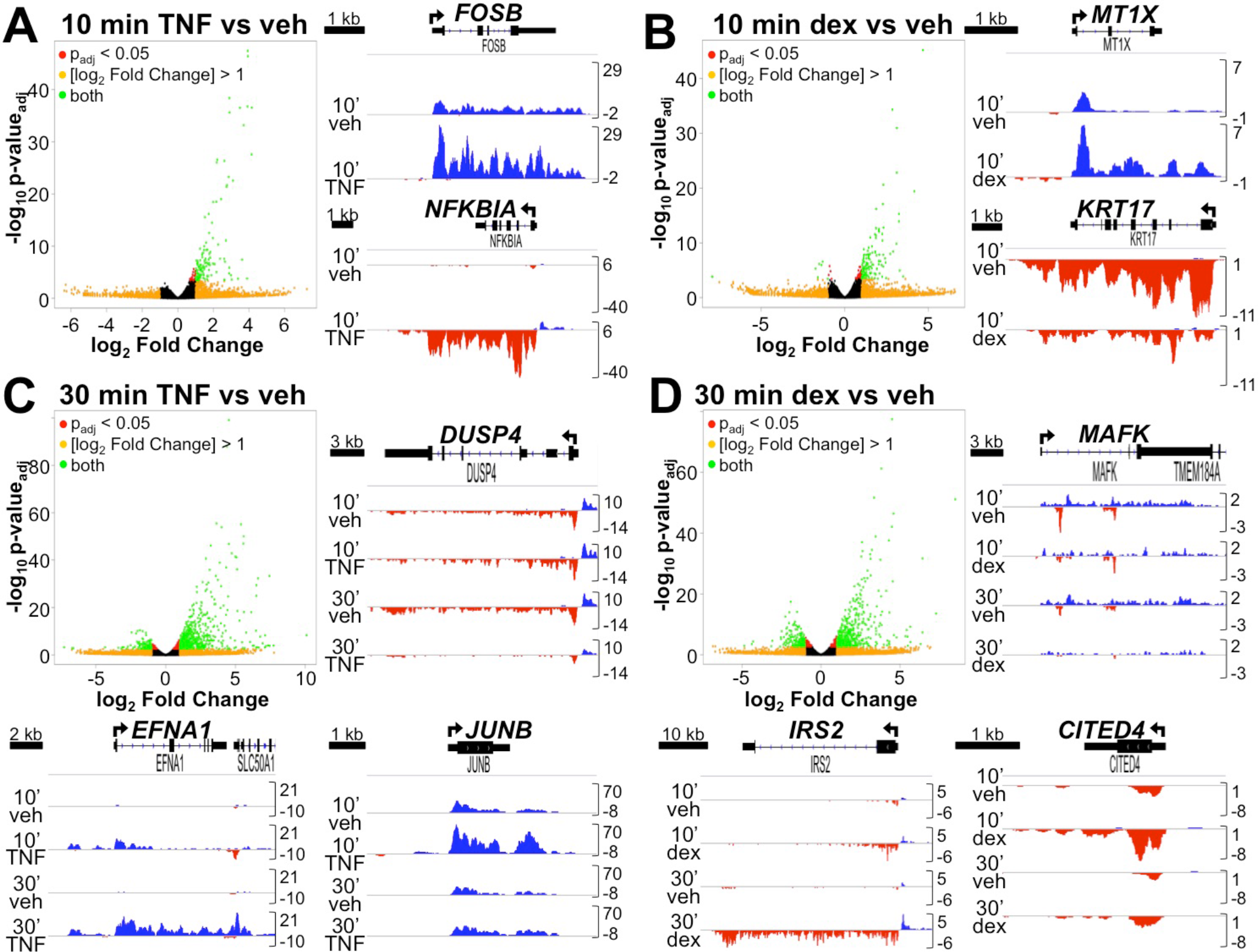
Primary transcriptional effects of TNF and glucocorticoids determined by GRO-seq. (A-D, left/top) Volcano plots indicating differentially regulated nascent transcripts identified by GRO-seq in Beas-2B cells treated with (A) 10 min TNF vs. veh, (B) 10 min dex vs. veh, (C) 30 min TNF vs. veh, and (D) 30 min dex vs. veh. Representative examples are shown to the right/bottom of each volcano plot as GRO-seq tracks visualized in the Integrative Genomics Viewer (IGV) browser based on counts per million mapped reads (vertical scales). Treatment conditions are indicated on the left of each track. Positive (blue) peaks are reads annotated to the positive/sense strand while negative (red) peaks reflect reads annotated to the negative/antisense strand. The TSS and direction of transcription are indicated by arrows at the top of each screenshot.

Comparison of the 10 and 30 minute time points revealed that the genome-wide response is more pronounced at 30 minutes for both TNF and dex treatment, with 995 and 783 genes differentially transcribed, respectively (Fig. 1C and D; the complete set of differentially transcribed genes are in Supplemental File S1). Glucocorticoid-mediated repression was also more readily detectable after 30 minutes of treatment with 237 genes and long non-coding RNAs (lncRNAs) showing demonstrably lower levels of transcription (e.g. *MAFK*, Fig. 1D), of which ~20% exhibited evidence of repression at 10 minutes based on non-adjusted p < 0.05. Because of the more robust transcriptional response, we selected the 30-minute time point for detailed analysis of the signaling pathway interactions between glucocorticoids and TNF, which were subsequently classified into three dominant clusters. In cluster one (Fig. 2A), comprising 117 genes, dex and TNF cooperated to enhance the transcription of target genes, a pattern that we have previously linked to anti-inflammatory effects of glucocorticoids (Kadiyala et al. 2016). In cluster two (Fig. 2B), comprising 75 loci, dex repressed the transcription of TNF target genes, whereas reciprocally, TNF exerted a repressive effect on dex-mediated induction of 28 transcripts in cluster three (Fig. 2C). Supporting this transcription-based mechanistic organization, gene ontology analysis of differentially transcribed genes reveals a unique set of functional terms for each cluster (Supplemental Table S1).

**Figure 2.**
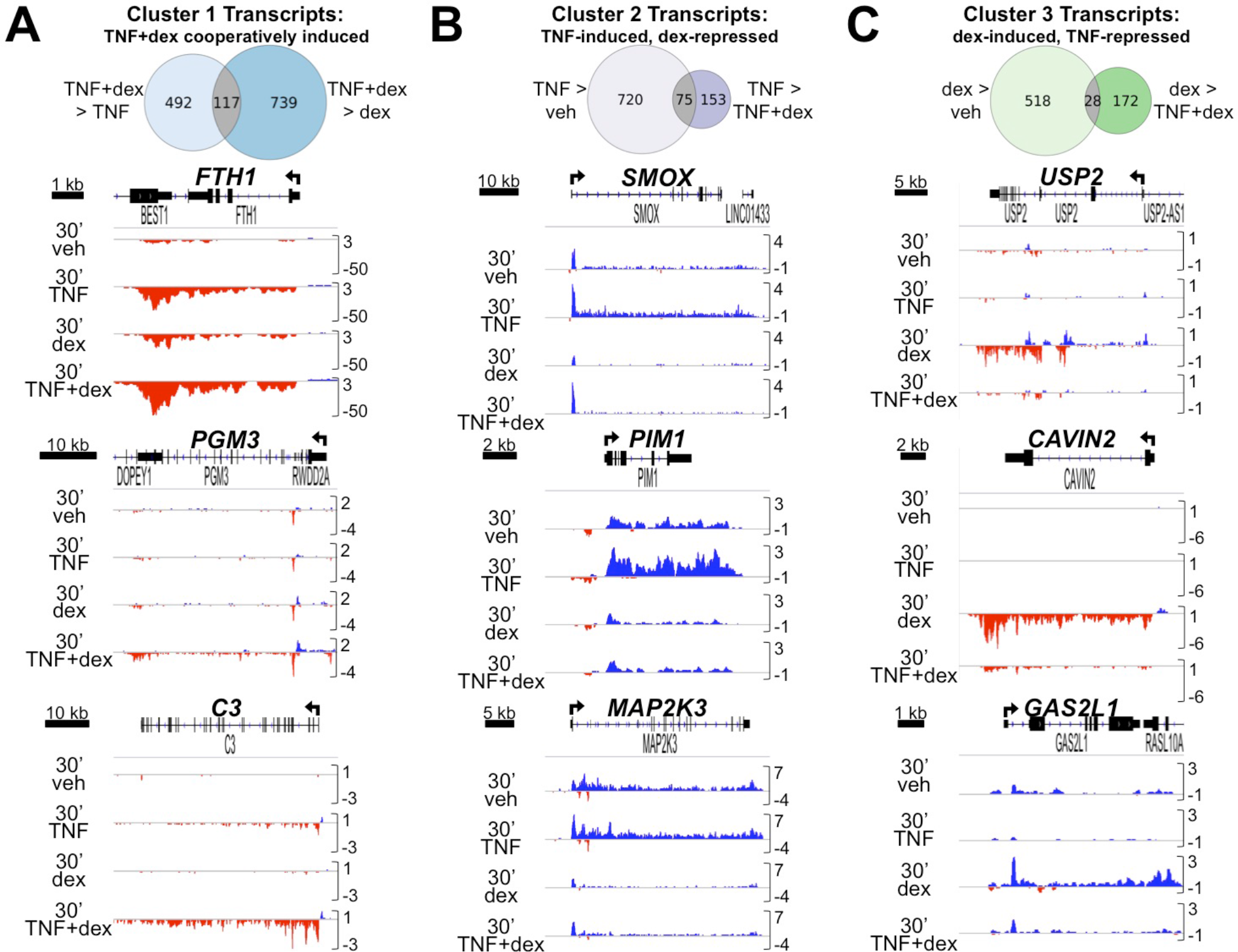
GRO-seq reveals distinct patterns of transcriptional crosstalk between TNF and dex. (A-C, top) Criteria used to cluster transcripts by indicated pattern of TNF+dex regulatory crosstalk and Venn diagrams showing how many differentially regulated transcripts (based on p_adj_ < 0.05) following 30 min treatment met the established criteria. (A-C, bottom) IGV screenshots of GRO-seq data, as described for Figure 1, illustrating examples of each crosstalk pattern.

### Glucocorticoid signaling rapidly represses basal enhancer activity

Previous studies have demonstrated that bidirectional transcription exclusive of gene transcription start sites, herein referred to as enhancer RNAs (eRNAs), can be used to characterize active enhancers (Allen et al. 2014; Azofeifa et al. 2018). We therefore used Transcription fit (Tfit), a machine learning program modeling RNAPII activity, to annotate putative eRNAs (Azofeifa and Dowell 2017). After merging putative eRNA regions across all samples, we obtained the set of active enhancers within this cell type (n=75,395) and subsequently utilized DESeq2 to assess differential RNAPII activity in response to TNF, dex or combinatorial treatment, as detailed in the Methods section; complete DESeq2 analysis is in Supplemental File S2. Not unexpectedly, the patterns of enhancer activity largely mirrored the underlying regulatory pattern of nearby gene-encoding transcripts. Thus, we were able to group enhancers into similar crosstalk patterns as we had defined for gene transcription (Fig. 3A).

**Figure 3.**
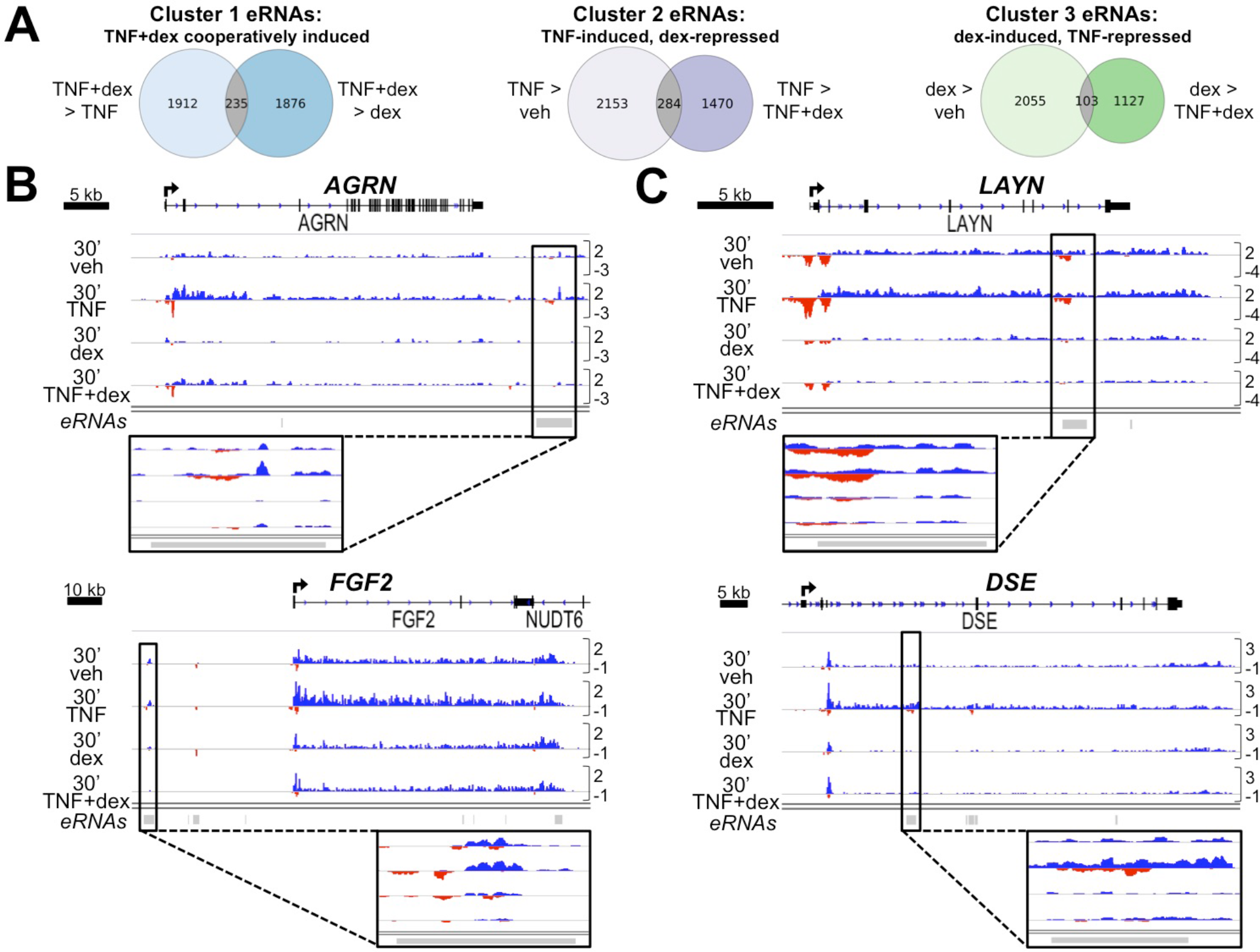
Glucocorticoids promptly repress basal activity of TNF-induced enhancers defined by bidirectional GRO-seq signature. (A) Criteria used to cluster bidirectional enhancer signatures by TNF+dex crosstalk pattern and Venn diagrams illustrating the number of differentially transcribed eRNAs (using non-adjusted p-value < 0.05) meeting these criteria. (B-C) IGV-visualized GRO-seq tracks of Cluster 2 (TNF-induced, dex-repressed) transcripts and nearby (B) intergenic or (C) intragenic called eRNAs (signified by gray bars at the bottom of each screenshot) exhibiting a similar regulatory pattern. Zoomed-in views of each relevant bidirectional signature are provided.

To further explore mechanisms of inflammatory repression by glucocorticoids, we specifically scrutinized enhancers that exhibited reduced activity based on GRO-seq data with TNF + dex treatment in comparison to TNF alone (analogous to cluster 2 in Fig. 2). Reduced enhancer activity with TNF + dex treatment versus TNF alone was, based on non-parametric testing, strongly associated (p < 0.00001) with lower absolute levels of enhancer activity after dex treatment relative to vehicle alone. Representative examples of dex-mediated repression of the basal activity of TNF-responsive enhancers and associated genes are shown in Fig. 3. As NF-kB exhibits minimal basal occupancy in Beas-2B cells (Kadiyala et al. 2016), these data do not support a singular mechanistic role for repressive tethering between GR and NF-kB as the basis for dex-mediated repression of TNF signaling.

We were interested in determining whether GRO-seq defined enhancer activity could, in an agnostic fashion, identify NF-kB and GR as the factors responsible for driving transcriptional effects of TNF and dex respectively. To accomplish this, we used motif displacement (MD) analysis (Azofeifa et al. 2018), a metric that calculates the enrichment of transcription factor motifs within a 150 bp radius relative to motif frequency within a 1500 bp radius centered on Tfit-called enhancers. Comparing vehicle to TNF, the MD score for p65 was the most increased across MD scores for all 641 transcription factors defined within the HOCOMOCO database (Supplemental Fig. S2) (Kulakovskiy et al. 2018). There was no significant change in the MD score for the GR motif between vehicle and dex treatment. These data indicate that NF-kB binding sites are statistically enriched at the center of TNF-regulated enhancers, whereas regulation of enhancer transcription by GR may be less correlated with central enrichment for the consensus GR motif, as defined within the HOCOMOCO database.

### ChIP-seq suggests glucocorticoids exert repressive effects on open chromatin without direct GR occupancy

To further assess the basis for primary repressive effects of dex on transcription, we analyzed dex-repressed enhancers defined by GRO-seq in the context of our previously published ChIP-seq data (Kadiyala et al. 2016). Intriguingly, many of the dex-repressed enhancers exhibited presumptive GR binding peaks under basal culture conditions (Supplemental Fig. S3), a phenomenon of unclear significance that we have noted previously (Kadiyala et al. 2016; Sasse et al. 2017), but that has also been reported in murine studies of the effects of exogenous glucocorticoids (Lim et al. 2015). These presumptive GR binding peaks were present at many of the same genomic locations, although with reduced intensity, after dex treatment. This occupancy pattern contrasts sharply with canonical GR binding regions (So et al. 2007), which exhibit minimal occupancy under basal conditions and robust peaks in the presence of dex. Possible explanations for the ChIP-seq peaks present at dex-repressed enhancers under basal culture conditions include a non-canonical interaction between GR and chromatin in the absence of supplemental ligand, or alternatively, these peaks may be a consequence of nonspecific interactions between certain GR antibodies and so-called hyper-ChIPable genomic regions, which are typically associated with open chromatin (Teytelman et al. 2013).

As an initial step to distinguish between these possibilities, we performed Western blots for GR protein on cytoplasmic and nuclear fractions of Beas-2B cells. We found no evidence of GR in the nucleus in the absence of dex treatment (Supplemental Fig. S4), suggesting that the GR ChIP-seq peaks seen in vehicle treatment likely represent nonspecific interactions between this GR antibody and chromatin. Moreover, many of the apparent GR binding peaks found under basal culture conditions overlap with regions that exhibit classic features of open chromatin (Tsompana and Buck 2014; Diehl and Boyle 2016; Castillo et al. 2017), including histone H3K27 acetylation, binding of numerous transcription factors, and DNase1 hypersensitivity, as observed in ENCODE datasets (see Supplemental Fig. S3). Thus, while our previously reported ChIP-seq data do not directly refute possible GR occupancy at dex-repressed enhancers, contextualization of these data strongly raises the possibility that well-described nonspecific interactions between antibodies and open chromatin may confound analysis of GR occupancy at these loci (Krebs et al. 2014; Jain et al. 2015).

To determine more definitively whether GR directly occupies dex-repressed enhancer regions, we performed additional GR ChIP-seq experiments in cells transduced with a lentiviral shRNA targeting GR (shGR), or with a control construct (shCtrl). We performed two independent experiments, each in duplicate, using two different antibodies: GR-lA1, the antibody we used previously that showed significant peaks under basal culture conditions, and a new antibody we generated, GR-356, which showed much lower occupancy levels under basal culture conditions in pilot experiments, despite being generated against the same epitope as GR-lA1. Both antibodies recognize a single protein that migrates at the predicted size of GR and is reduced in cells expressing the shGR construct (Supplemental Fig. S5). To align with our prior ChIP-seq studies, Beas-2B cells were treated for one hour with vehicle, TNF, dex, or TNF + dex. As expected, well-established canonical sites of GR occupancy (e.g. the *FKBP5* and *PDK4* loci, Fig. 4A) exhibited minimal ChIP-seq peaks under basal culture conditions, but robust peaks after dex treatment. Binding at these sites was abrogated in shGR cells. These results at specific regulatory elements were assessed genome-wide through MD analysis on the ChIP-seq datasets from both antibodies (Azofeifa et al. 2018). When applying the MD score approach to ChIP, we centered on MACS-called ChIP-seq peaks within a dataset. This revealed highly significant central enrichment for the canonical dimeric GR binding motif in shCtrl cells treated with dex (MD scores of 0.43 and 0.67 with GR-lA1 and GR-356, respectively). Consistent with ~90% GR knockdown achieved in cells transduced with shGR (Supplemental Fig. S5), enrichment for the GR motif was significantly abrogated (p < 0.00005), but not wholly eliminated, in ChIP-seq data from cells transduced with shGR (Fig. 4B and Supplemental Fig. S6). In contrast, MD scores for two potential matches to the so-called nGRE motif ranged from 0.13 to 0.15. These scores are similar to the expected score of 0.1 for a motif that is neither enriched nor excluded from the center of a ChIP-seq peak set, i.e. a randomly distributed motif. Moreover, these scores were not significantly altered in cells transduced with shGR (Fig. 4B and Supplemental Fig. S7), rendering it extremely unlikely that GR interacts specifically with either of these nGRE sequences. Taken together, as anticipated, ChIP-seq data generated with either GR antibody demonstrate highly significant genome-wide association between GR and canonical dimeric binding motifs.

**Figure 4.**
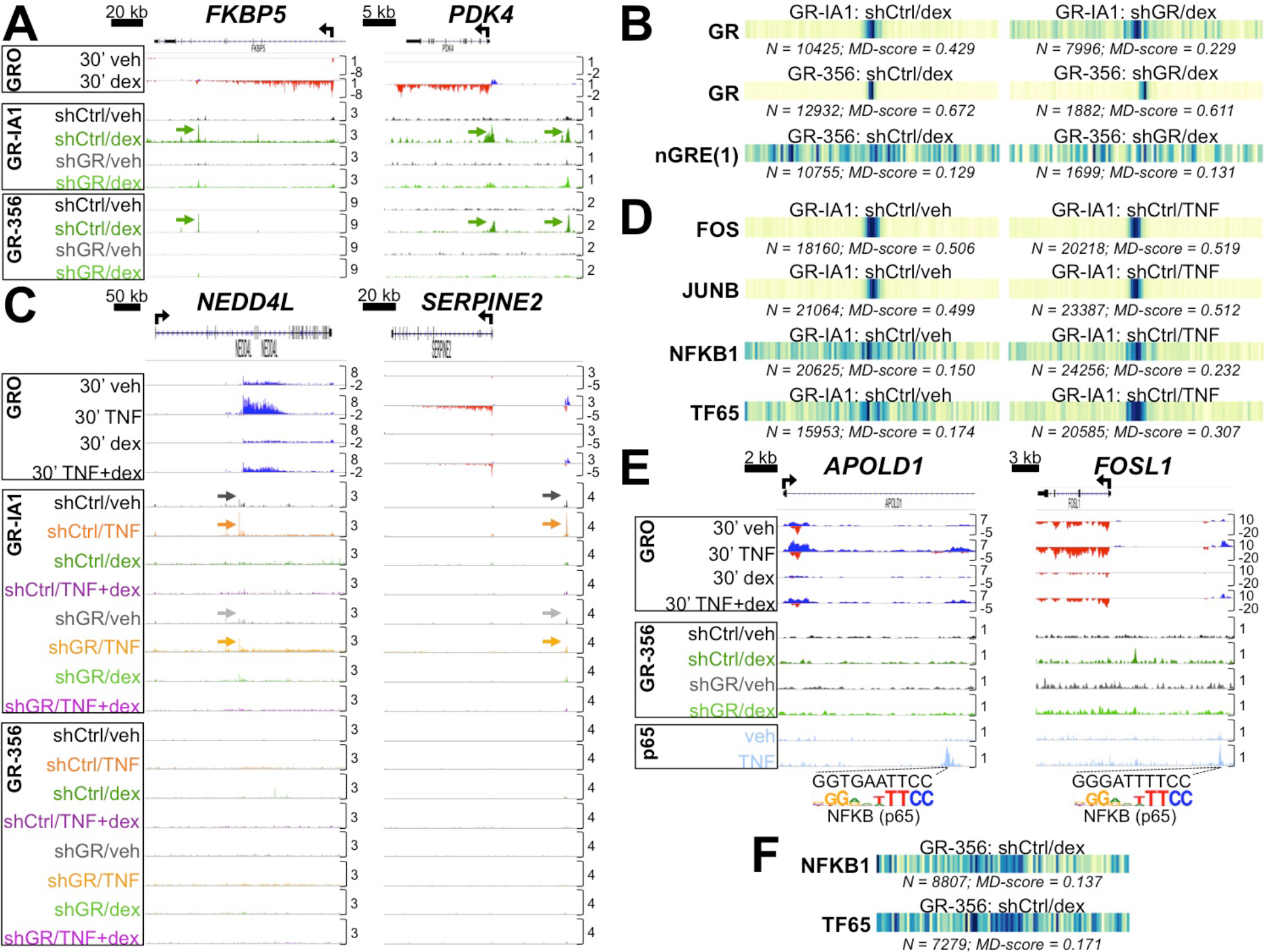
GR ChIP-seq with stable GR knockdown excludes tethering transrepression and direct enhancer occupancy by GR as major mechanisms underlying primary glucocorticoid repression of TNF targets. (A) Aligned GRO-seq and GR ChIP-seq IGV tracks at representative dex-induced loci; arrows indicate GR peaks associated with canonical GR binding sites. Vertical scales indicate maximum counts per million mapped. (B) Motif displacement (MD) analysis depicting frequency of overlap between GR and nGRE binding motif matches within ±1500 bp of GR ChIP-seq peak summits in the indicated datasets; darker colors signify greater enrichment. (C) ChIP-seq (and GRO-seq) IGV tracks at dex-repressed loci exhibiting apparent GR occupancy in veh-and TNF-treated shCtrl cells (indicated by arrows) with the GR-IA1 but not the GR-356 antibody. (D) MD analysis of enrichment for indicated NFKB and AP-1 family binding motifs in select GR-IA1 ChIP-seq datasets. (E) GR-356 ChIP-seq (and GRO-seq) data combined with p65 ChIP-seq tracks (Kadiyala et al. 2016) at Cluster 2 loci illustrating dex-mediated repression of p65-occupied enhancers containing NFKB/p65 binding site matches in the absence of detectable GR occupancy. (F) MD analysis for indicated NFKB family binding motifs in select GR-356 ChIP-seq datasets.

Unlike the concordant results from both antibodies indicative of classically reported GR signaling and occupancy at canonical binding sites, ChIP-seq occupancy patterns in vehicle-and TNF-treated cells showed substantial differences between the two antibodies. Specifically, for the lA1 antibody, despite a lack of nuclear GR, MACS-identified peaks were observed in basal culture conditions (n=68,278 peaks), including at TNF-induced enhancers (Fig. 4C), in association with highly significant MD scores for AP-1 family members (Fig. 4D). Minimal enrichment for canonical NF-kB binding motifs was also detected, which was significantly increased with TNF treatment (Fig. 4D), despite the absence of nuclear GR in TNF-stimulated cells (Supplemental Fig. S4). The MD scores for AP-1 and NF-kB family members were not significantly reduced in shGR cells in comparison to shCtrl (Supplemental Fig. S6 and S7). Taken together, given the lack of nuclear GR and the failure of stable GR knockdown to consistently reduce the MD scores for AP-1 and NF-kB motifs, these data indicate that the lA1 ChIP-seq peaks present in vehicle and TNF treated cells do not represent *bona fide* interactions between the GR protein and chromatin. Instead they represent the occurrence of non-specific interactions between this GR antibody and these hyper-ChIPable genomic regions, which are strongly enriched for AP-1 and, to a lesser extent, NF-kB binding motifs.

In contrast to the lA1 antibody, the 356 ChIP-seq data showed minimal detectable MACS-identified peaks with vehicle and TNF treatment (n=251 and 271 peaks, respectively; e.g. Fig. 4C), suggesting that the 356 antibody is less prone to artifactual interactions with chromatin. Thus, to determine whether local GR occupancy is required for repression of TNF-regulated enhancers, we integrated the GRO-seq data, the 356 ChIP-seq results and our previously published ChIP-seq data for the p65 subunit of NF-kB (Kadiyala et al. 2016). Enhancers with inducible p65 occupancy that were repressed by dex without evidence of GR occupancy were readily identified in these integrated data (Fig. 4E). Thus, tethering interactions between GR and NF-kB and/or direct occupancy of GR at specific TNF-induced enhancers is not a requirement for dex-mediated repression of their activity. Supporting this notion, MD scores ranged between 0.14 and 0.17 for NF-kB binding motifs within the dex-treated 356 ChIP-seq dataset (Fig. 4F), indicative of only minimal central enrichment for NF-kB family binding sequences. Moreover, the MD scores for NF-kB family members are lower than MD scores for motifs associated with numerous other transcription factor families, including AP-1 (top score 0.46), BACH (top score 0.31), RUNX (top score 0.29), NFI (top score 0.28), TEAD (top score 0.24), and MAF (top score 0.23) families, amongst others (Supplemental Fig. S7; the entire set of MD scores are available in Supplemental File 3). These data are consistent with well-described combinatorial transcriptional regulation by GR and other transcription factors (Diamond et al. 1990; Wang et al. 1999), which is nucleated by DNA binding sites for GR and these factors, rather than a protein-tethering regime that would encompass interactions between GR and such widely diverse transcription factors. In that regard, aligned with our prior report of very high affinity GR binding sites within a subset of p65 occupied regions (Kadiyala et al. 2016), we detected juxtaposition of NF-kB binding motifs and canonical GR binding motifs within our GR-356 ChIP-seq data. Specifically, within ~13000 GR ChIP-seq peaks located within a 1500 bp radius of a canonical GR motif, there were 1882 matches for the NF-kB motif. GR-p65 co-regulatory interactions through canonical binding sites for each factor can result in cooperative gene induction (e.g. Fig. 2A) as we have previously described, but can also be associated with repression (Luecke and Yamamoto 2005; Uhlenhaut et al. 2013). In either case, in aggregate our data indicate that primary repressive effects of dex on the expression of TNF target genes can occur without evidence of local GR occupancy within the enhancer element, and are not generally mediated through either nGREs or tethering interactions between GR and NF-kB.

### Glucocorticoids cause rapid changes in chromatin accessibility

It was intriguing that the pattern of non-specific interactions between the lA1 antibody and chromatin changed dynamically, manifested by increases in the MD scores for NF-kB family members in the lA1 ChIP-seq datasets from TNF- versus vehicle-treated cells (see Fig. 4D). Scores for AP-1 family members also decreased with the addition of dex in comparison to vehicle treatment (Supplemental Fig. S7). Although no single architectural feature is entirely predictive of hyper-ChIPable chromatin regions (Li et al. 2016), the changes in these non-specific interactions potentially indicate that dex treatment directly modifies the chromatin structure of these enhancers. To test directly the hypothesis that glucocorticoid treatment results in rapid changes in basal chromatin structure at TNF-induced enhancers, we performed high-resolution micrococcal nuclease (MNase) accessibility assays on representative dex-repressed enhancers in comparison to dex-induced enhancers, all of which exhibited regulation by dex treatment in the GRO-seq data. Accordingly, assays were performed in Beas-2B cells treated with vehicle or dex for 10, 30, or 60 minutes. Using tiled qPCR, we imputed relative protection from MNase digestion, a reflection of chromatin accessibility (Infante et al. 2012), across ~700-1500 bp of each enhancer. These data show a rapid increase in protection from MNase digestion within part of each GR-repressed enhancer (*PTPRK* and *TMEM217* in Fig. 5A-B; *SMURF2* in Supplemental Fig. S8); notably these enhancers did not exhibit significant occupancy with the GR-356 antibody under any treatment condition (Supplemental Fig. S9). In contrast, protection from MNase cleavage was reduced within the two dex-induced enhancers tested with this assay (*ANGPTL4* and *SH3RF3* in Fig. 5C-D), and these enhancers showed significant dex-induced GR-occupancy (Supplemental Fig. S9). For each of our interrogated enhancers, the flanks were protected from digestion, consistent with positioned nucleosomes flanking these regulatory elements. Thus, rapid intranucleosomal changes in enhancer chromatin structure can occur with dex treatment, characterized by decreased access to MNase digestion at dex-repressed enhancers and increased access to MNase digestion at dex-induced enhancers.

**Figure 5.**
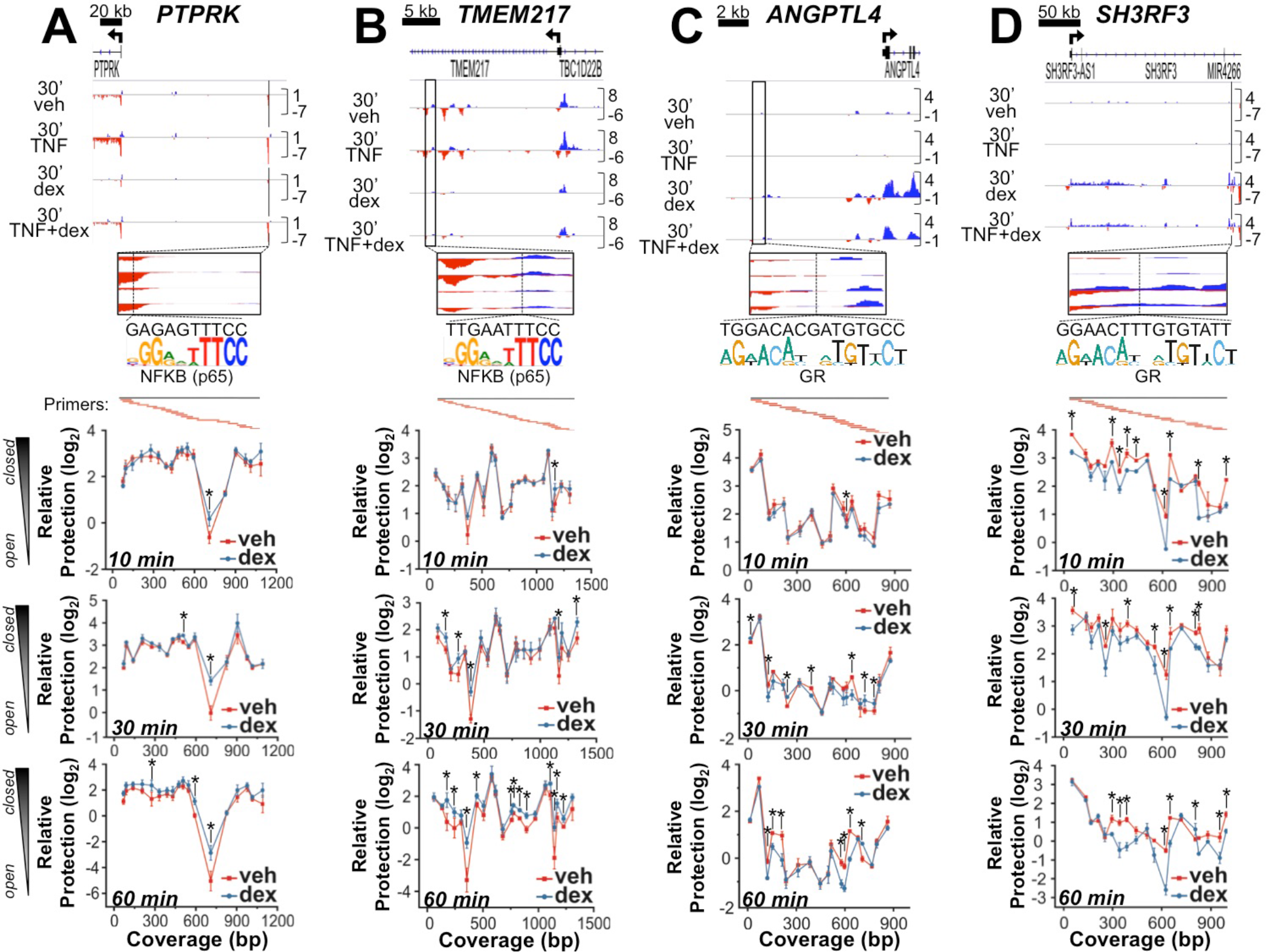
Dex treatment rapidly changes basal chromatin structure of both TNF- and dex-induced enhancers. (A-D, top) IGV screenshots of GRO-seq data with solid black lines/rectangles showing specific TNF-induced (A-B) and dex-induced (C-D) enhancer regions that were interrogated by the MNase assay. Below each screenshot is a zoomed-in view of each assayed region, including the location (dotted black line) and sequence of the strongest match to the NFKB/p65 consensus binding motif (A-B) and GR binding motif (C-D). Beneath the binding site matches are the locations of overlapping tiled qPCR primers (amplicons in red) that span each region and correspond to the data points in the line graphs below. (A-D, bottom) Mean relative protection (±SD) against MNase cleavage of each target region as measured by qPCR in Beas-2B cells treated with veh or dex for 10, 30, or 60 min (*p < 0.05 vs. veh). Greater protection indicates less accessibility or a more closed chromatin structure, as illustrated on the far left.

### Reporter analysis confirms enhancer activity and suggests a chromatin requirement for rapid repression

The above data suggest that local changes in chromatin structure are associated with rapid dex-mediated repression, despite no evidence of direct GR occupancy at these enhancers. To determine whether native chromatin structure and genomic context are a requirement for glucocorticoids to exert rapid repressive effects, we cloned putative enhancers associated with genes that exhibited different patterns of TNF-dex interactions as defined by GRO-seq and interrogated their activity in reporter assays. For this analysis, we exploited the “NanoLuc” system, a destabilized, highly luminescent short half-life shrimp luciferase reporter that can parallel endogenous increases and decreases in eukaryotic gene expression at 60 minute resolution (Masser et al. 2016). Using this assay system, enhancers from cluster one (i.e. cooperative-regulation by TNF and dex) mimicked the behavior of the endogenous enhancer and showed cooperative regulation by TNF and dex after just 60 minutes of treatment (Fig. 6A). In contrast, repressive effects of dex on cluster two enhancers were not evident at the one hour time point (Fig. 6B), and in the case of *IER3*, were not detected until 4 hours. Similar to cluster one, enhancers from dex-induced genes (cluster three) also exhibited increases in activity with dex after one hour of treatment (Fig. 6C). Thus, inductive effects of dex, including cooperation between dex and TNF, are recapitulated by isolated enhancer fragments, whereas rapid repressive effects of dex on enhancer activity depend at least in part on chromatin and/or genomic context.

**Figure 6.**
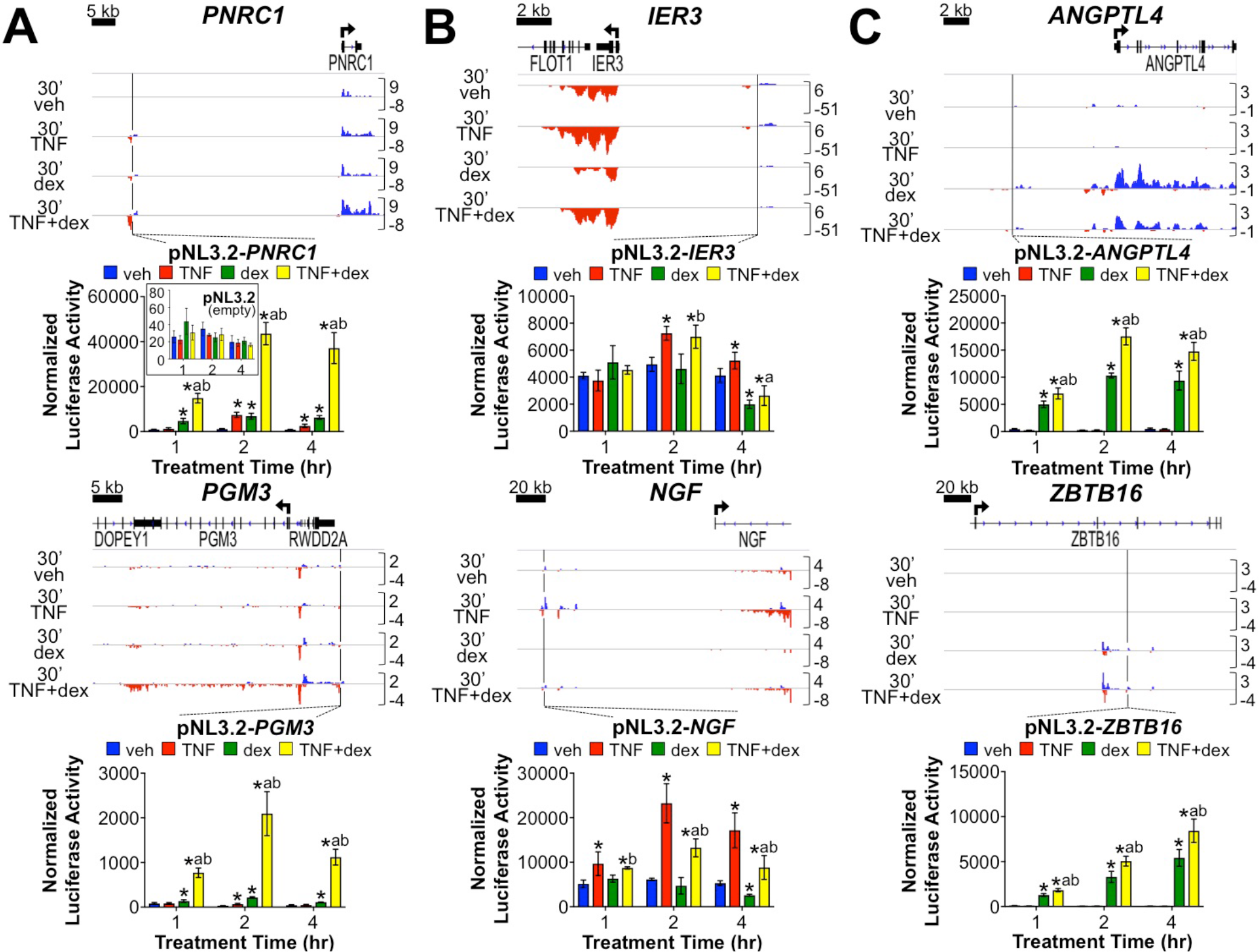
Enhancer reporter assays suggest chromatin requirement for immediate early repression of TNF-induced enhancer activity by glucocorticoids. (A-C) GRO-seq tracks and bar graphs showing relative genomic locations and normalized luciferase activity, respectively, of (A) Cluster 1 (TNF+dex cooperatively induced), (B) Cluster 2 (TNF-induced, dex-repressed) and (C) Cluster 3 (dex-induced, TNF-repressed) enhancers cloned into a destabilized, short half-life luciferase reporter vector and transiently transfected into Beas-2B cells prior to treatment with vehicle, TNF, dex, or TNF+dex for 1, 2, or 4 hours. Luciferase activity of empty vector control (pNL3.2) is included in the inset of the pNL3.2-*PNRC1* graph in Panel A. Data from two representative reporters are shown in each panel. Bars indicate mean activity (+SD) normalized to that of a firefly luciferase internal control. *p < 0.05 vs. veh; ^a^p < 0.05 vs. TNF; ^b^p < 0.05 vs. dex.

## DISCUSSION

In this study, we set out to determine whether glucocorticoids exert primary repressive effects on gene expression, defined as repression occurring without a protein intermediate. Our GRO-seq data, which show reduced nascent transcription in some genomic regions after just 10 minutes of dex treatment, provide strong evidence in support of direct repressive effects. Repression of many TNF-regulated enhancers and genes occurred in the absence of TNF treatment, indicating, as suggested by others (Jubb et al. 2016; Oh et al. 2017), that tethering between GR and p65 is not required for a primary response. Moreover, the previously described nGRE sequence was not significantly enriched within GR occupied regions, indicating that widespread repression does not occur through direct interactions between GR and nGREs in our system. Instead, our analysis of GR ChIP-seq data with and without stable GR knockdown suggests that many dex-repressed enhancers reside in so-called hyper-ChIPable regions of the genome that are subject to non-specific interactions with antibodies in ChIP assays. Dynamic changes in these non-specific interactions between GR antibodies and enhancer chromatin with dex and TNF treatment, in conjunction with MNase accessibility assays of representative enhancers, indicate that dex treatment rapidly reduces access to chromatin at repressed enhancers. Taken together, our data establish that glucocorticoids exert primary repressive effects on transcription through altering chromatin structure. However, the two dominant explanations for primary repression, GR tethering to NF-kB and interactions between GR and nGREs, are unable to account for these effects.

What is the mechanistic basis for primary GR-mediated repression? Although we cannot entirely rule out the existence of direct interactions between GR and repressed enhancers that lead to reduced enhancer activity, our data provide little support for this notion and clearly refute an important role for GR tethering repressively to NF-kB. Instead, our data support a model in which dex-induced genome-wide binding of GR to canonical binding sites results in reciprocal tightening of chromatin structure at selected enhancers that are not directly occupied by GR (Fig. 7). Although speculative, this could occur through a variety of mechanisms including allosteric or biophysical effects of GR-nucleated transcription complexes on chromatin topology, three-dimensional interactions between GR and the repressed enhancers, and/or redistribution of co-regulators that are required to maintain open chromatin structure at the dex-repressed enhancers. Conversely, proteins that tend to repress transcription, such as H1 histones (Hergeth and Schneider 2015), could potentially be liberated from GR-induced regulatory regions and subsequently establish more compact chromatin structures at other genomic regions (Weintraub 1984). Irrespective of the mechanistic basis for the primary repressive effects of glucocorticoids, our observation that many GR-repressed enhancers are in genomic regions with hyper-ChIPable characteristics could explain why dynamic GR occupancy at repressed enhancers has been variably reported (Lim et al. 2015), thus providing parsimony for seemingly discordant data.

**Figure 7.**
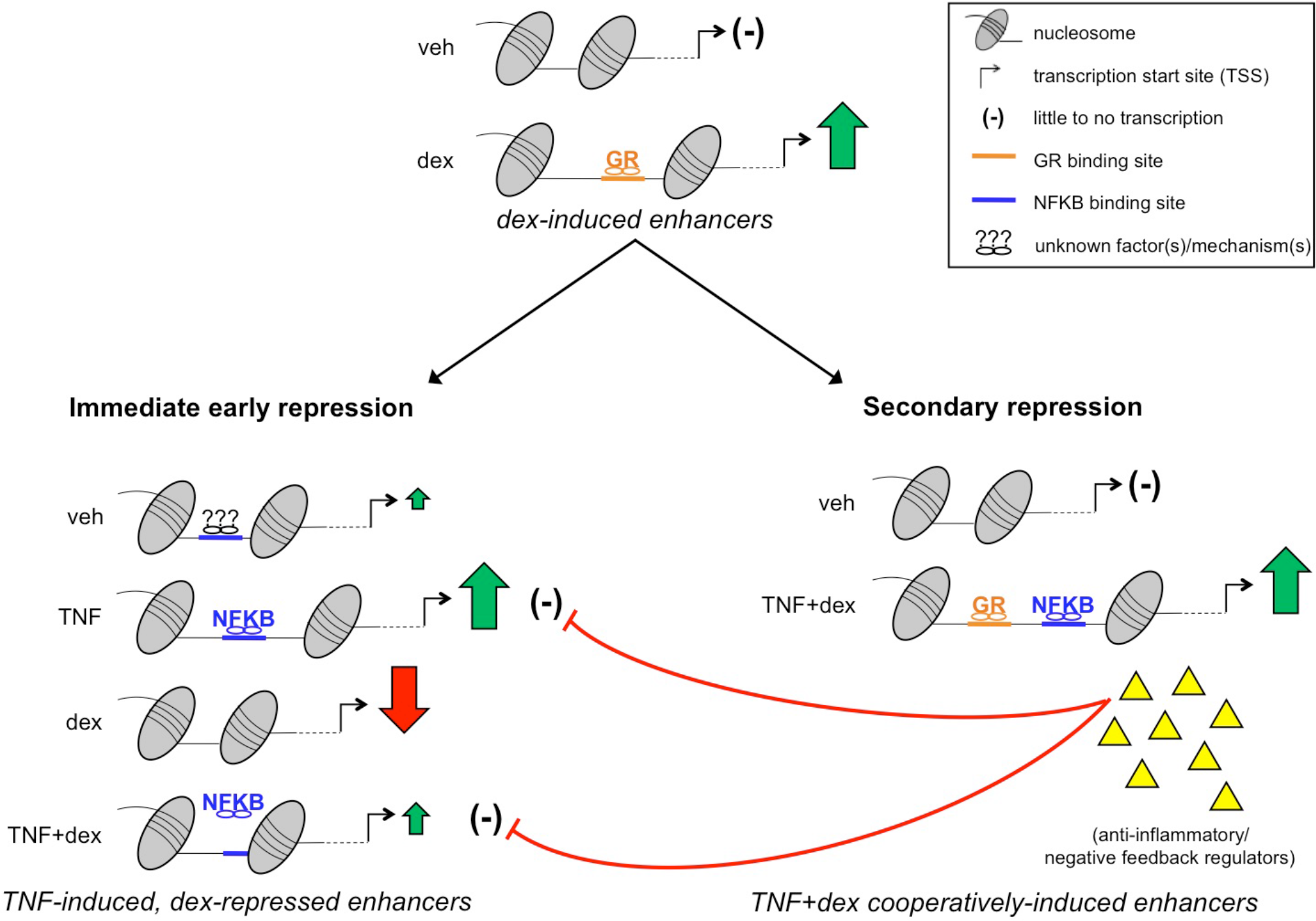
Two-step model of glucocorticoid-mediated inflammatory repression. Glucocorticoids rapidly induce target genes predominantly through glucocorticoid receptor (GR) interactions with enhancers harboring canonical GR binding sites (GBSs). The consequences of genome-wide transcriptional induction by GR include (1) an immediate early wave of repression, characterized by rapid reciprocal intranucleosomal tightening at select TNF-induced enhancers that is independent of local GR occupancy, and (2) a secondary wave of repression involving interactions with NFKB at specialized enhancers that requires local GR occupancy to cooperatively induce transcription/translation of remarkably potent anti-inflammatory effectors. These effectors exert robust negative feedback control to limit further NFKB activity and also function to counteract/destroy the various pro-inflammatory products (e.g. cytokines, chemokines, proteases) generated during the initial response, thereby playing a crucial role in promoting complete resolution of inflammation.

It has been argued that redistribution of limiting amounts of specific co-regulators from one set of enhancers to another is a unifying explanation for rapid repression in association with transcriptional induction (Guertin et al. 2014; Schmidt et al. 2016), which has been observed in a variety of other signaling contexts (Hah et al. 2011; Step et al. 2014; Franco et al. 2015; Loft et al. 2015; Toropainen et al. 2016). Although redistribution of FOXA1 occupancy with glucocorticoid treatment has been reported (Swinstead et al. 2016), whether specific transcription factors or co-activators rapidly redistribute their nuclear position from repressed to activated regulatory elements in association with GR activation has not been fully investigated. Studies of transcription factor redistribution in association with transcriptional induction will, however, need to carefully account for the hyper-ChIPable nature of regulatory regions that are subject to the rapid repressive and chromatin remodeling effects of glucocorticoids. Indeed, whereas numerous studies have reported on chromatin structure remodeling in association with GR signaling (Voss et al. 2011; Jubb et al. 2017; McDowell et al. 2018), our MNase data indicate that remodeling at some sites is associated with changes in intranucleosomal accessibility (e.g. *PTPRK*, Fig. 5A), rather than broad changes in nucleosome positioning. This change in intranucleosomal access may lead to dynamic changes in artifactual interactions between antibodies and these chromatin regions being observed in ChIP-seq datasets, as we found here in our studies with the GR-IA1 antibody. Co-factor redistribution, or “squelching” models may also need to be broadened to include local effects of chromatin and nuclear positioning, as our data indicate that timely repression of some enhancers depends on chromatin context. Whether primary repression in association with intranucleosomal chromatin tightening is a property of physiologic nuclear receptor signaling also remains to be determined. In that regard, it is intriguing to speculate that normal circadian pulses of corticosteroids, which are known to reduce cytokine expression (Gibbs et al. 2014), may restore “chromostasis” or a neutral chromatin structure in cells that have experienced activation of inflammatory genes during the course of the preceding 24 hours.

In addition to demonstrating that glucocorticoid signaling exerts primary repressive effects on the transcription of inflammatory genes, our GRO-seq data build on prior work from our group and others reporting on cooperative glucocorticoid–TNF crosstalk resulting in secondary repression of inflammatory genes (Vettorazzi et al. 2015; Kadiyala et al. 2016).

Specifically, the high resolution afforded by GRO-seq allowed us to discover additional target genes in which dex + TNF treatment resulted in higher levels of RNAPII activity than was evident with dex or TNF alone. Novel targets of glucocorticoid-TNF cooperation include *PGM3*, whose deficiency is associated with elevated IgE levels and asthma (Yang et al. 2014); *FTH1*, which protects against TNF-mediated apoptosis (Pham et al. 2004); *C3*, a traditionally pro-inflammatory gene that was recently reported to suppress stress-associated apoptosis in Beas-2B airway epithelial cells (Kulkarni et al. 2018); and *ZFAND5*, a TNFAIP3-like inhibitor of NF-kB (Huang et al. 2004). The anti-inflammatory functions of each of these genes further implicate cooperative gene regulation by GR and NF-kB as an important mechanistic underpinning of repression of inflammation and cell injury by glucocorticoids.

A number of classically inductive signaling pathways, including estrogen signaling, exert primary repressive effects on transcription that encompass repression of TNF target genes (Ruan et al. 2003; Franco et al. 2015). The estrogen receptor can also cooperate with NF-kB to induce gene expression (Franco et al. 2015). Why then do glucocorticoids function as uniquely potent anti-inflammatory drugs? The data we have presented here, in combination with prior work reporting on cooperative anti-inflammatory gene regulation by GR and NF-kB (Altonsy et al. 2014; Vettorazzi et al. 2015; Sasse et al. 2016), provide insight into this unmatched clinical efficacy. Specifically, our data support a two-step model (see Fig. 7) in which inductive gene regulation by GR results in rapid reciprocal repression of inflammatory enhancers through a chromatin-based repressive mechanism. Whereas this form of reciprocal repression is not exclusive to glucocorticoids, through canonical binding sites for GR and NF-kB, the activity of a subset of specialized enhancers controlling anti-inflammatory genes is uniquely augmented by glucocorticoids in cooperation with NF-kB. This GR-specific regulatory system renders these anti-inflammatory genes relatively resistant to both primary repression and to negative feedback control of NF-kB, thus decoupling the expression and resulting biologic effects of these negative feedback regulators from pro-inflammatory cytokine expression. With respect to the airway, the range of genes we have found that are regulated through this cooperative mechanism is remarkable and encompasses genes that protect against various forms of cellular stress, as well as powerful negative feedback regulators of NF-kB, including TNFAIP3. Our data thus provide an explanation for the clinical effectiveness of glucocorticoids in reversing inflammation associated with asthma and chronic obstructive pulmonary disease exacerbations, which are typically a consequence of airway infections that both induce NF-kB signaling and cause cellular injury (Kersul et al. 2011; Hoppenot et al. 2015; Nicod and Kolls 2015; Schuliga 2015). Our findings may also enable rational pharmacologic improvement of glucocorticoid-mediated inflammatory repression, a long sought goal in pulmonary therapeutics (Barnes 2006; Clark and Belvisi 2012).

## METHODS

### Cell culture, reagents and Western blotting

Beas-2B cells (ATCC) were cultured in Dulbecco’s Modified Eagle Medium (DMEM; Corning) containing L-glutamine and 4.5 g/L glucose and supplemented with 10% fetal bovine serum (VWR) and 1% penicillin/streptomycin (Corning). Dexamethasone (dex; Sigma) was dissolved in sterile 100% ethanol (vehicle) and used at a final concentration of 100 nM. Recombinant Human TNF-alpha Protein purchased from R&D Systems was diluted in 1X Dulbecco’s phosphate buffered saline (DPBS) containing 0.1% bovine serum albumin (BSA) and used at a final concentration of 20 ng/ml. Primary antibodies for Western blotting included mouse anti-TNFAIP3 [59A426] (ab13597), rabbit anti-NFKBIA [E130] (ab32518) and rabbit anti-Beta-Tubulin (ab52901) from Abcam, anti-GAPDH [FL-335] (sc-25778) from Santa Cruz Biotechnology, rabbit anti-CEBPB (PA5-27244) from Thermo Fisher Scientific, and anti-Lamin B1 [D4Q4Z] (12586S) from Cell Signaling Technology. Secondary antibodies were ECL sheep anti-mouse IgG-HRP (95017-332) and donkey anti-rabbit IgG-HRP (95017-330) from GE/Amersham. Western blotting and protein detection were performed using standard procedures (Sasse et al. 2016). Antibodies for GR ChIP-seq were GR-IA1 (Sasse et al. 2017) and GR-356, which was raised in rabbit against the identical epitope as IA1 (QPDLSKAVSLSMGLYMGETETKVMGNDLG) and prepared by Covance. The shGR construct was cloned as described (Kampmann et al. 2014) and targeted the following sequence: TGGTGTCACTGTTGGAGGTTAT.

### Global Run-on Sequencing (GRO-seq)

Our GRO-seq protocol was based on previous publications (Allen et al. 2014) and is detailed in the Supplemental Methods. Data has been deposited in GEO (GSE124916).

### GRO-seq computational analysis

#### Data processing, visualization and identification of eRNAs

Two biological replicates of each treatment (veh, TNF, dex, and TNF + dex) at both 10- and 30-minute timepoints were processed using a standardized nascent transcription data pipeline. The random priming kit used to prepare GRO-seq libraries returns reads in antisense (3’ to 5’) orientation, so reads were first converted to their reverse complement (5’ to 3’) using FASTX Toolkit fastx reverse complement (v. 0.0.13) with the argument ‘−Q33’. Reads were trimmed for adapters, minimum length, and minimum quality using the BBMap Suite (v. 38.05) bbduk.sh tool with arguments ‘ref = truseq.fa.gz literal = CCCGTGTTGAGTCAAATTAAGCCGCAGGCTCCACTCCTGGTGGTGCCCTT ktrim = r qtrim = 10 k = 23 mink = 11 hdist = 1 maq = 10 minlen = 2’. FastQC (v. 0.11.5) was performed and calculated a median Illumina phred quality score of 40 both pre-and post-trimming, which indicates an inferred base call accuracy greater than 99.9% for all bases in all samples. Trimmed reads were mapped to the human genome (hg38 downloaded from Illumina iGenomes on August 8th, 2018, with corresponding Bowtie2 index files) using Bowtie2 (v. 2.1.0) in ‘–end-to-end’ alignment mode with ‘–very-sensitive’ preset options, which resulted in a median of 92.48% of reads being aligned for all samples. A complete report for all quality control metrics and mapping results can be viewed in Supplemental File S4, which was generated using MultiQC (v. 1.6). Conversion of SAM files to BAM format, and subsequent generation of bedgraph files and visualization in the Integrative Genomics Viewer (IGV; v. 2.4.10) is described in the supplementary methods. Output from application of FStitch and Tfit to identify regions with bidirectional transcriptional activity is in Supplementary File S5.

#### Differential transcription analysis of genes and bidirectionals/enhancers

Using the RefSeq : NCBI Reference Sequences for hg38, including both NM and NR accession types (downloaded from the UCSC track browser on May 18, 2018), counts were calculated for each sorted BAM file using multibBamCov in the BEDTools suite (v. 2.25.0). Genes (NM accession type) and lncRNAs (NR accession type) were then filtered such that only the isoform with the highest number of reads per annotated length was kept in order to minimize duplicate samples being included in differential transcription analysis. DESeq2 (v. 1.20.0, Bioconductor release v. 3.7) was then used to determine which genes were differentially transcribed between the different treatments for each time point separately (Supplemental File S1). For bidirectional/enhancer comparisons, all bidirectional prediction Tfit calls were first merged using mergeBed (argument −d 60) from the BEDTools suite (v. 2.25.0) to generate an annotation file. Counts were then calculated for each sample using multicov from the BEDTools suite (v. 2.25.0) and DESeq2 was used to calculate differentially transcribed bidirectionals/enhancers (Supplemental File S2). Supplemental scripts and data processing information are available at https:/github.com/Dowell-Lab/Sasse2019.

### Lentiviral production and transduction of Beas-2B cells

HEK293 FT cells were grown to ~70% confluence on Poly-D-Lysine-coated 10 cm tissue culture dishes. Cells were transfected with lentiviral packaging vectors pMDLG/RRE, pMD2.G and pRSV/Rev (2.88 ug total, Addgene) plus pMK1221-control or −GR shRNA (2.88 ug shCtrl or shGR, respectively) using TransIT-293 Transfection Reagent (Mirus Bio) as instructed by the manufacturer. Following 72 hr incubation, viral supernatant was collected from HEK cells, centrifuged at 1000 rpm for 10 min at room temperature and filtered with a sterile 0.45 um syringe filter. Beas-2B cells were plated on 10 cm tissue culture dishes and grown to confluence. Viral transduction was performed using a 1:1 ratio of viral supernatant to standard growth media along with 8 ug/ml polybrene. Cells were incubated for 24 hr and then transduced a second time, followed by antibiotic selection with puromycin (1 ug/ml) for 24-48 hr.

### Chromatin immunoprecipitation-sequencing (ChIP-seq)

Beas-2B cells transduced with lentiviral shCtrl or shGR were grown to confluence in 10 cm tissue culture dishes (~10E6 cells/plate) and treated with vehicle, TNF, dex, or TNF+dex for 1 hour. Cells were cross-linked by adding 16% methanol-free formaldehyde to a final concentration of 1% and incubating for 5 min at room temperature. ChIP was then performed as described (Sasse et al. 2013), with the exception that samples were sonicated on high power for 35 cycles of 30-sec bursts separated by 30-sec incubations in ice water with a Diagenode Bioruptor. Samples were immunoprecipitated with 12 ug of GR-356 or GR-IA1 antibody. One ng of purified ChIP DNA was used to prepare uniquely barcoded libraries with the Ovation Ultralow Library System from NuGEN. Libraries were pooled and sequenced in duplicate on an Illumina NovaSEQ using 2 × 150 bp paired-end reads by the Microarray and Genomics Core at the University of Colorado-Denver. Computational analysis of ChIP-seq data is described in detail in the Supplemental Methods.

### Micrococcal nuclease (MNase) chromatin accessibility assay

MNase chromatin accessibility assays were largely performed as described (Infante et al. 2012). Beas-2B cells were grown to confluence in 10 cm tissue culture dishes and treated with vehicle or dex (100 nM) for 10, 30, or 60 min. Crosslinking was performed by adding 16% methanol-free formaldehyde directly to the culture media to a final concentration of 1% and incubating for 5 min at room temperature. Formaldehyde was quenched by addition of glycine to a final concentration of 125 mM and incubation for 10 min at 4°C. Cells were washed in ice-cold 1X DPBS for 5 min at 4°C and then scraped off dishes in ice-cold immunoprecipitation lysis buffer (50 mM HEPES-KOH pH 7.4, 1 mM EDTA, 150 mM NaCl, 10% glycerol, 0.5% Triton X-100) supplemented with 1X Protease Inhibitor Cocktail (Thermo Scientific) and nutated for 30 min at 4°C, followed by dounce homogenization (50 strokes). Nuclei were collected by centrifugation at 600 x g for 5 min at 4°C and resuspended in ice-cold nuclease digestion buffer (10 mM Tris-HCl pH 8.0, 1 mM CaCl_2_). MNase (10 Units/sample; New England Biolabs) was added and reactions were incubated for 40 min at 37°C. Digestion was stopped by addition of an 8.6% SDS/7 mM EDTA solution and crosslinks were reversed by adding Proteinase K (20 mg/ml; Life Technologies) and incubating 6 hr at 65°C. DNA was then extracted using Phenol:Chloroform:Isoamyl Alcohol (25:24:1; Sigma), precipitated in 100% ethanol, washed in 70% ethanol, air-dried for 30 min, then resuspended in 1X TE Buffer. DNA was separated on a 2% agarose gel and bands between 120-150 bp were excised and purified (QIAquick Gel Extraction Kit, Qiagen). Chromatin accessibility was assayed using quantitative RT-PCR (qRT-PCR) as described (Sasse et al. 2013). Primers (80-120 bp) were designed using an overlapping (40-60 bp overlap between adjacent primers) head-to-tail tiling method to completely span each enhancer region of interest. Primer efficiencies from genomic DNA amplification were used to normalize experimental Ct values. Assays were generally performed in biologic quadruplicate and repeated at least three times with qualitatively similar results. Primer sequences used for the MNase assays are listed in the Supplementary Table 2.

### Plasmids, transfection and reporter assays

Genomic enhancer regions of interest identified by GRO-seq were PCR-amplified, inserted into pCR2.1-TOPO (Life Technologies), then introduced into the pNL3.2 [NlucP/minP] minimal promoter-driven NanoLuc luciferase expression vector from Promega. Primer sequences for cloning, cloning strategies and transfection assay methods are described in detail in the Supplementary Methods.

### DATA ACCESS

GRO-seq data have been deposited at GEO (GSE124916). ChIP-seq data have been deposited at GEO (pending).

## ACKNOWLEDGMENTS

Work supported in part through NIH R01HL109557 (SKS, ANG); NIH R01GM125871 (MAG, RDD). ANG thanks Stephen J. Tapscott and his lab for insightful comments on ChIP artifacts.

## DISCLOSURE DECLARATION

RDD is a founder of Arpeggio Biosciences. All other authors have nothing to declare.

